# CRISPR-Cas targeting in *Haloferax volcanii* promotes within-species gene exchange by triggering homologous recombination

**DOI:** 10.1101/2025.06.18.660351

**Authors:** Deepak Kumar Choudhary, Israela Turgeman-Grott, Shachar Robinzon, Uri Gophna

## Abstract

CRISPR-Cas systems provide adaptive immunity in bacteria and archaea against mobile genetic elements, but the role they play in gene exchange and speciation remains unclear. Here, we investigated how CRISPR-Cas targeting affects mating and gene exchange in the halophilic archaeon *Haloferax volcanii*. Surprisingly, we found that CRISPR-Cas targeting significantly increased mating efficiency between members of the same species, in contrast to its previously documented role in reducing inter-species mating. This enhanced mating efficiency was dependent on the Cas3 nuclease/helicase and extended beyond the targeted genomic regions. Further analysis revealed that CRISPR-Cas targeting promoted biased recombination in favour of the targeting strain during mating, resulting in an increased proportion of recombinant progeny that are positive for CRISPR-Cas. To test whether an increase in recombination is sufficient to increase mating efficiency, we tested whether strains lacking the MRE11-RAD50 complex, which are known to have elevated recombination activity, also exhibited higher mating success. Indeed, these strains showed higher mating, as did cells that were exposed to DNA damage using methyl methanesulfonate. These findings suggest that CRISPR-Cas systems may contribute to speciation by facilitating within-species gene exchange while limiting between-species genetic transfer, thereby maintaining species boundaries.

## Introduction

CRISPR (Clustered Regularly Interspaced Short Palindromic Repeats)-Cas (CRISPR-associated genes) systems are now recognized as a predominant anti-viral defense that provides bacteria and archaea with adaptive immunity against mobile genetic elements (MGEs)^1,2^ during infection, CRISPR-Cas systems target foreign DNA and acquire short sequences, called spacers, from invading genomes. These spacers are integrated into the CRISPR locus, serving as immune memory for targeting and preventing future infections^3^. CRISPR loci consist of short repetitive elements (repeats) interspersed with spacers, along with a set of *cas* (CRISPR-associated) genes^3,4^. Cas proteins serve multiple functions: some act as effector endonucleases, others acquire new spacers, and some are responsible for crRNA maturation^4,5^.

Halophilic archaea use a unique cell fusion-based mating process to exchange chromosomal and plasmid DNA with and between species^6,7^. After the initial mating event, heterozygous cells are initially formed, containing genetic material from both parents. This stage can later lead to the formation of two different types of progeny: recombinant cells, with chromosomal loci from both parental strains, and cells that may segregate and revert to the original (parental) genotypes^7^. To study mating frequency, selection for mating products is applied such that heterozygous cells will be able to grow, as well as recombinants with a favourable combination that contains the markers that are selected for, while cells that revert to the parental genotypes will be unable to grow.

Previous mating experiments between different species of the halophilic archeon *Haloferax* have shown that when the CRISPR-Cas systems of one species targets that of another, the efficiency of productive mating events is reduced compared to non-targeting mating^8^. This raised the possibility that CRISPR-mediated degradation may cause mating cells to detach from their “offensive” mating partner prematurely, reducing the opportunity for gene exchange. Conversely, CRISPR-Cas mediated genome cleavage could actually stimulate homologous recombination (HR) during mating, because due to activity of the Cas3 nuclease/helicase of type I-B CRISPR-Cas systems (the most common system in halophilic archaea^5,9^. The latter is much more likely to occur when HR is efficient due to perfectly identical homology arms, as is often the case within species^6,10^.

In this study, we tested whether within-species CRISPR-Cas targeting affects mating success. Our results demonstrate that CRISPR-Cas targeting increases the frequency of gene exchange by influencing the mating process. Furthermore, we show that CRISPR-Cas targeting biased the recombination in favour of the targeting strain during mating, which could benefit CRISPR-Cas containing strains of Haloferax species, in which CRISPR-Cas is plasmid-encoded rather than ancestral. Importantly, CRISPR-Cas targeting can increase gene exchange within-species, while reducing between-species exchange, making these systems a factor that contributes to speciation.

### Within-species mating is increased by CRISPR-Cas targeting

Halophilic archaea can form cytoplasmic bridges that facilitate cell-to-cell contact, enabling genetic material transfer through a process called mating, which promotes horizontal gene transfer (HGT)^7,11,12^. Our lab previously demonstrated that CRISPR-Cas targeting reduces inter-species mating between *H. volcanii* and a very genetically distant species *H. mediterranei*, thereby limiting HGT^8^. To further understand CRISPR-Cas role in gene exchange among halophilic archaea, we investigated its effect on within-species mating.

We inserted a 40 bp spacer sequence into the *H. volcanii* genome. This sequence is targeted by a type I-B CRISPR-Cas system spacer that mediates DNA cleavage, known as interference^5,8,13^ (Figure 1A). We then introduced this sequence into a strain lacking CRISPR-Cas genes to prevent self-targeting and autoimmunity. Next, we conducted mating experiments using WR532 (*ΔpyrE2*) that has the wild type CRISPR-Cas system and two *H. volcanii* strains, either UG633 (*ΔhdrB, ΔtrpA, Δcas* genes, containing the target spacer inside *ΔtrpA*) or UG634 (*ΔhdrB, ΔtrpA, cas+*) as a non-targeted control strain (Figure 1A). Surprisingly, in six out of seven biological replicates, within-species CRISPR-Cas targeting significantly **increased** mating efficiency compared to the non-targeted control strain (Figure 1A & B).

**Figure 1.**
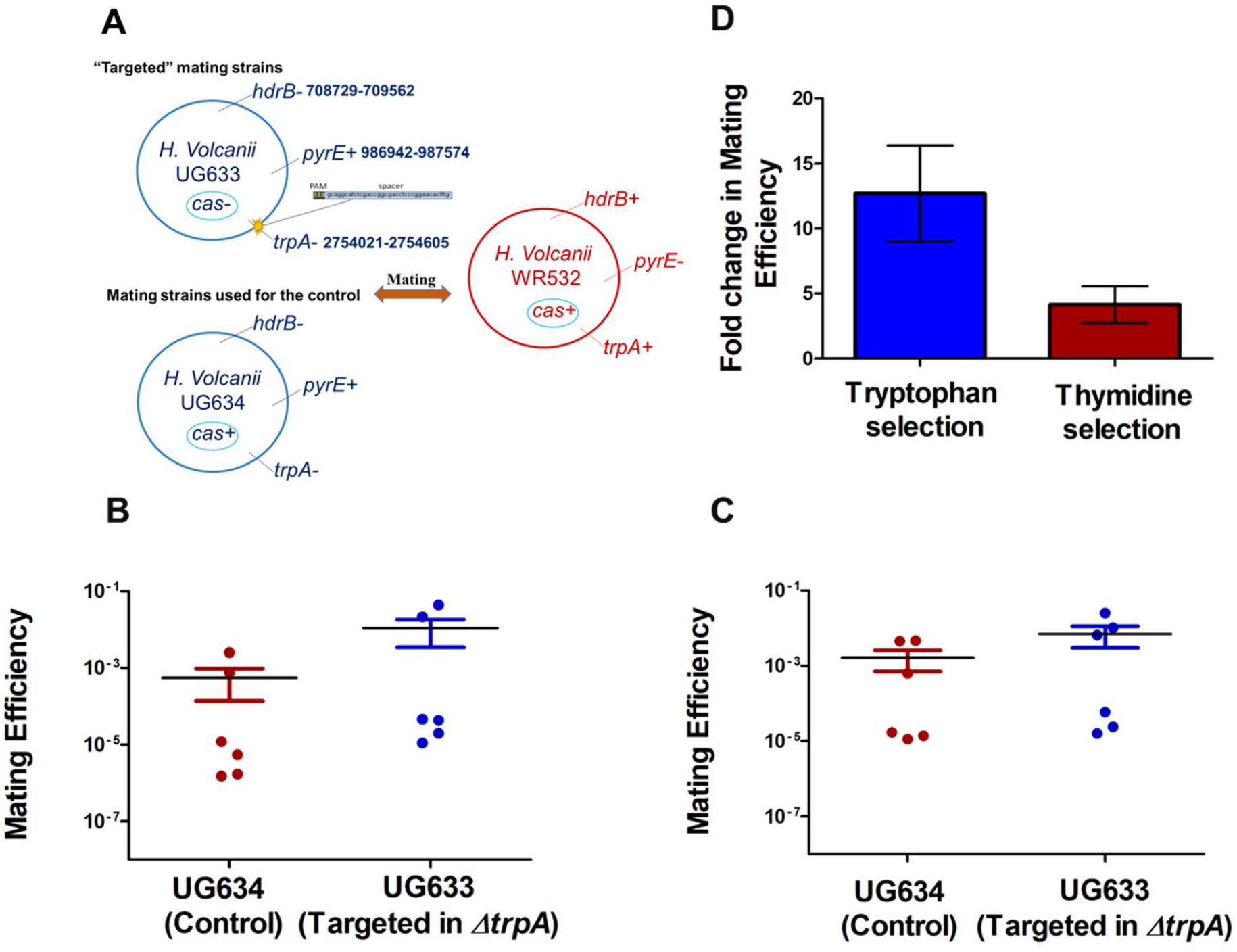
A, B, C & D. Within-species mating is increased by CRISPR-Cas targeting. **A**. Strains used for the mating experiments. The “targeted” strain was engineered to contain a validated *H. volcanii* CRISPR spacer+PAM (40bp)^13^ sequence, so that *H. volcanii* CRISPR-Cas could target it during mating. The location of the spacer+PAM sequence insertion is marked with a yellow color and corresponds to the locus of the deleted *trpA* gene. The mating experiments were conducted between UG633 (*ΔhdrB, ΔtrpA, Δcas genes*, which also contains the target of the spacer inside *ΔtrpA*) and WR532 (*ΔpyrE*). The UG634 (*ΔhdrB, ΔtrpA*) non-targeted strain was used as control. Following mating, the cells were plated on Hv-Ca+thymidine, and Hv-Ca+ tryptophan media. **B**. Mating results after selection for tryptophan. **C**. Mating results after selection for thymidine. The Wilcoxon matched-pairs signed-rank test was performed to compare targeted and non-targeted mating for both Figure A and Figure B, with p-values of P < 0.0331 and P < 0.0313, respectively **D**. Comparison of mating efficiency between experiments using thymidine vs. tryptophan selection.

Further, we investigated whether the increased mating efficiency was specifically due to CRISPR-Cas targeting rather than the CRISPR-Cas machinery itself, previously shown to affect recombination in *Haloferax* in the absence of targeting^9,14^. We conducted another mating experiment using two of the previously described strains: UG633 (*ΔhdrB, ΔtrpA, Δcas* genes, containing the target of the spacer inside *ΔtrpA*) and WR532 (*ΔpyrE2*) with an active CRISPR-Cas system: for this experiment, we used *H. volcanii* UG444 (*ΔhdrB, Δcas*) as a non-targeted control strain lacking the CRISPR-Cas system (Supplementary Fig. 1a). Consistent with our previous findings, in seven out of nine biological replicates, CRISPR-Cas targeting **increased** mating efficiency compared to the non-targeted control strain (Supplementary Fig. 1b). These results suggest that the increased mating efficiency was dependent on CRISPR-Cas targeting.

### CRISPR-Cas targeting affects gene exchange in loci distant from the targeted site

To test whether the effect of CRISPR-Cas targeting on mating success extends beyond the targeted region, we selected for a distant genetic marker by plating the mating culture on a medium that selects for presence of the *hdrB+* allele (using a medium containing *trpA+* and lacking *hdrB+*). The *hdrB* gene region is 801881 bp distant from the targeted *trpA* site. Mating experiments were conducted between UG633 (*ΔhdrB, ΔtrpA, Δcas* genes, with the target spacer inserted in *ΔtrpA*) and WR532 (*ΔpyrE2*) as shown in figure1A. After mating, we first plated the culture on a medium containing thymidine and lacking tryptophan to select for cells containing the *trpA+* marker. Interestingly, when we selected for tryptophan marker, we observed high mating efficiency in CRISPR-Cas targeting compared to the non-targeted control (Figure 1B).

To test whether the effect of CRISPR-Cas targeting extends beyond the targeted region, we plated the culture on a medium that selects for the *hdrB+* marker (using a medium containing tryptophan but lacks thymidine). **Surprisingly**, CRISPR-Cas targeting still resulted in high mating efficiency compared to the non-targeted control, although this efficiency was **approximately half** of what was observed with the *trpA-*based selection (Figure 1C &D). The increased mating efficiency was not limited to the region directly affected by CRISPR-Cas targeting (*trpA+*), suggesting that CRISPR-Cas targeting may globally enhance gene exchange.

### Cas3 is required for increased mating in the targeted strain

Cas3 is one of the most prominent CRISPR-associated proteins found in all Type I CRISPR-Cas systems, where it exhibits both helicase and nuclease activity. Upon the invasion of foreign DNA, Cas3 is recruited by Cascade and cleaves spacer-matching DNA sequences through its endonuclease activity^5,15,16^. We tested whether Cas3 is required for the increased mating observed under targeted conditions. Specifically, if the increased mating is due to CRISPR-Cas interference, then deleting Cas3 in a mating partner with a CRISPR-Cas system that interferes with the target should reverse the increased mating phenotype.

To test this, we chose a strain with a *cas3* deletion (but all other *cas* genes remain intact) and conducted a mating experiment between two sets of strains: UG633 *(ΔhdrB, ΔtrpA, Δcas* genes, which also contains the target of the spacer within *ΔtrpA*) and UG610 (*ΔpyrE, Δcas3*). A non-targeted strain, UG444 (*ΔhdrB, Δcas*), was used as a control (Figure 2A). After mating, the cells were plated on Hv-Ca plates, and unlike the previous experiment, no difference in mating was observed in the targeted strain UG633, compared with the non-targeted control (Figure 2B). These findings suggest that the increased mating observed under targeted conditions is due to CRISPR-Cas-mediated interference and is dependent on Cas3 proteins.

**Figure 2:**
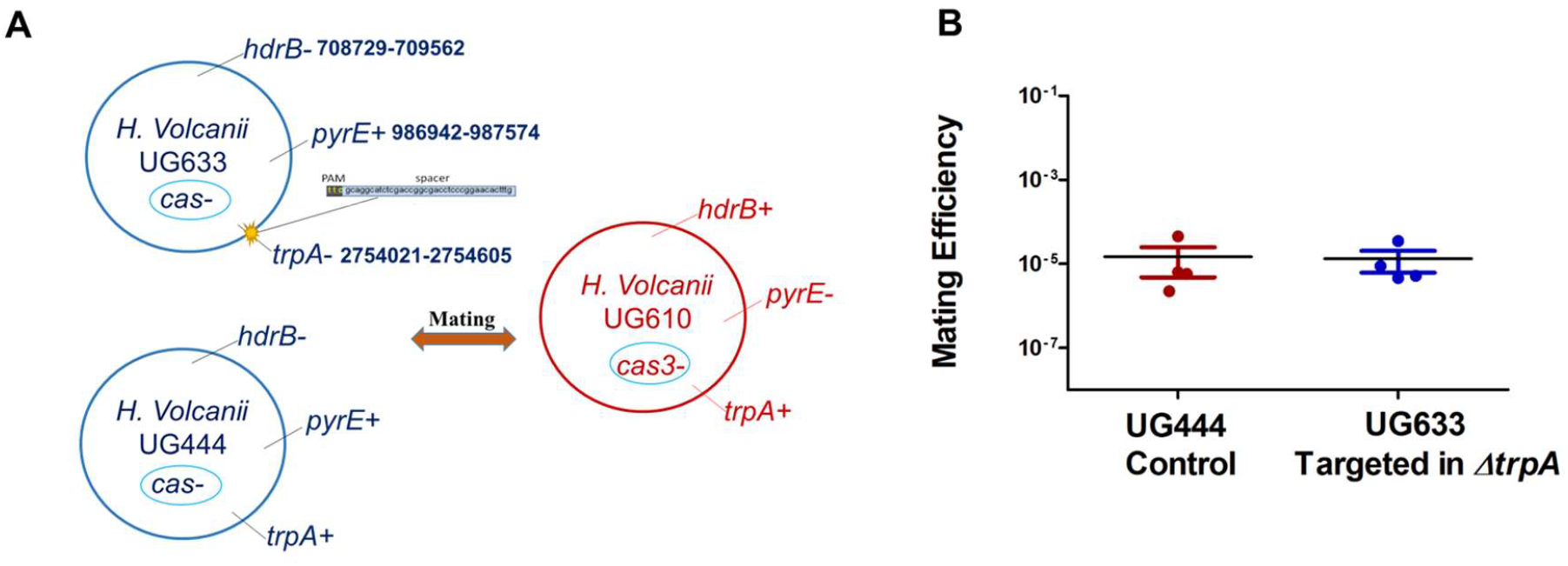
CRISPR-Cas activity is required for increased mating in the targeted strain. **A**. The mating experiment has been conducted between UG633 (*ΔhdrB, ΔtrpA, Δcas genes*, also contains the target of the spacer inside *ΔtrpA*) and UG610 (*ΔpyrE, Δcas3)*. UG444 (*ΔhdrB, Δcas*) non-targeted strain has been used for the control. **B**. Following mating, the cells were plated on Hv-Ca media.

### CRISPR-Cas systems drive biased recombination in favour of the targeting strain

CRISPR-induced genome cleavage has long been known to promote recombination between homologous chromosome arms and initiate homologous recombination (HR) in genome editing applications^17,18^. As we have observed that CRISPR-Cas targeting increases within-species mating efficiency, it is plausible that this change in mating efficiency could be a result of an increased recombination rate. This is because presumably cells can enter a mating state and then this state can be disrupted, but if recombination has already taken place, segregation will still result in the marker combination that is later selected for. However, CRISPR targeting may bias recombination because one mating partner cuts the genome of the other.

To determine whether CRISPR targeting indeed biases recombination, we took 30-50 colonies (mating products) of the experiments described above (Figures 1A & B), which were selected for the thymidine marker gene (from Hv-Ca +tryptophan plates). Screening by PCR for the trpA gene which was not selected for, indicated that all mating products were recombinant rather than heterozygous, since either the trpA-band or the trpA+ band were obtained but never both (supplementary figure S2). These recombinant mating products can be either WR532 cells that obtained the *pyrE* gene or UG633 cells that obtained the *hdrB* gene. We then performed replica plating onto Hv-Ca plates without additional supplementation on which only the former cells can grow because they are trpA (the trpA gene is located 801881 bp away from the *hdrB* gene so for the UG633 to recombine both is far less likely). Approximately 80% of colonies from the targeted mating experiment grew on Hv-Ca plates, whereas only 30%of colonies from the control plate grew on Hv-Ca plates across all biological replicates (Figure 3A and B), indicating that CRISPR targeting biases recombination, and favours gene acquisition by WR532.

**Figure 3.**
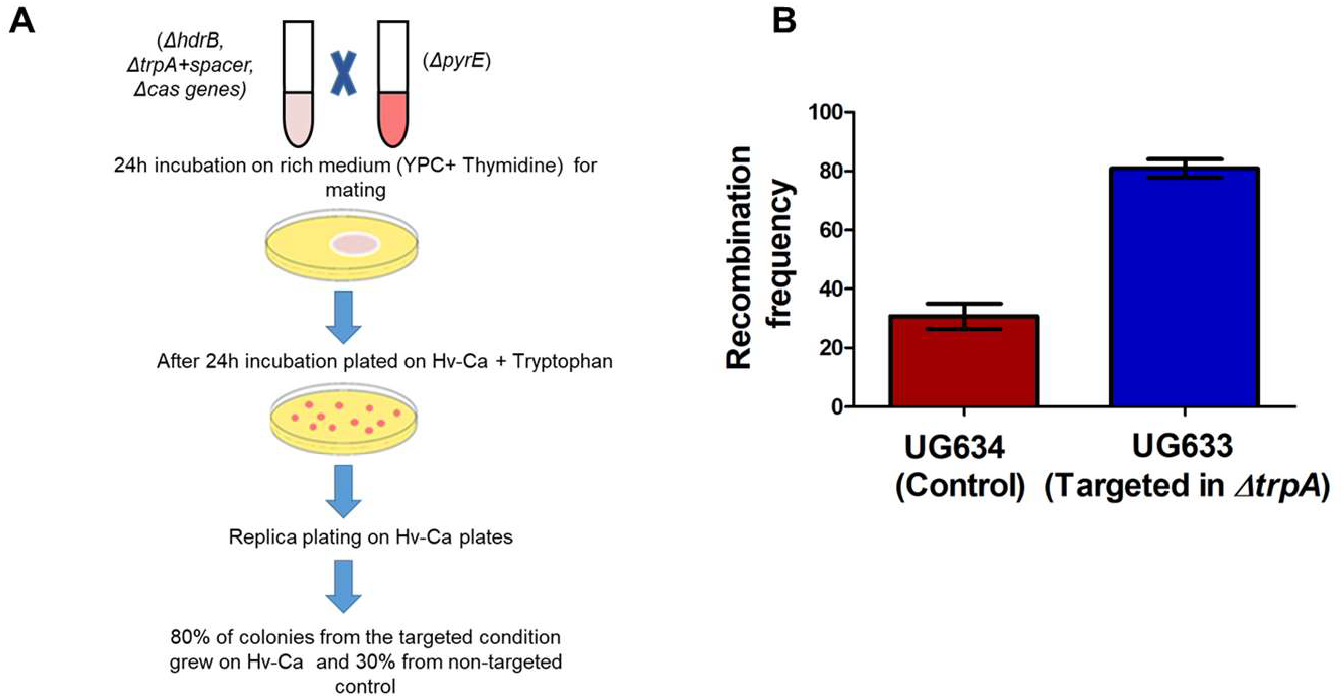
Recombination frequency is increased by CRISPR-Cas targeting. **A**. Schematic of recombination frequency. **B** Recombination frequencies by CRISPR-Cas targeting in within-species mating. Recombination frequencies were calculated by replica-plating on un-supplemented Hv-Ca media and based on the PCR results, as described above in the result. The results shown are an average of four independent experiments. Paired t-test was performed to compare targeted and non-targeted mating with p-values of P < 0. 0037.

Furthermore, when we took 50 colonies from the plate selected for the tryptophan marker from Hv-Ca +Thymidine plates (containing the target spacer inserted into *ΔtrpA*) and performed replica plating on Hv-Ca plates, bias was even stronger and, all colonies grew on the Hv-Ca plates, probably because UG633 cells whose genome was cut by CRISPR-Cas in the trpA locus failed to acquire the trpA gene from WR532 cells. We conclude that CRISPR targeting between strains results in a genetic antagonism where the mating partner being can provide genetic material to the “cutter” much more than vice versa, therefore benefitting strains that have CRISPR-Cas function.

### Disruption of *MRE11* and *RAD50* leads to increased mating efficiency in *H. volcanii*

A likely explanation for the increase in mating success is an increase in recombination. However, this effect of increased HR on mating success has never been examined. We therefore tested whether an *mre11-rad50* knock-out strain, known to have higher homologous recombination activity than the wild type^9,19^, will also show higher mating efficiency. We compared the mating efficiency of strain UG634 (Δ*hdrB*, Δ*trpA*) with either strain UG60 (Δ*mre11*, Δ*rad50*, Δ*pyrE*) or strain UG532 (Δ*pyrE*) that has wild-type *mre11*and *rad50*. As expected, we observed higher mating efficiency when the mating partner was deleted for *mre11* and *rad50*. This supports the central role of homologous recombination in determining archaeal mating success (Figure 4A). We were then curious to investigate whether inducing double-strand DNA breaks using methyl methanesulfonate (MMS) could promote recombination in lab strains of haloarchaea. To address this hypothesis, we conducted a mating experiment (cultures were incubated during fusion stage on YPC+MMS plates and YPC plates as a control for 24 hours) comparing strains with and without previous exposure to MMS. Our results revealed that MMS exposure increased mating events (Figure 4B).

**Figure 4.**
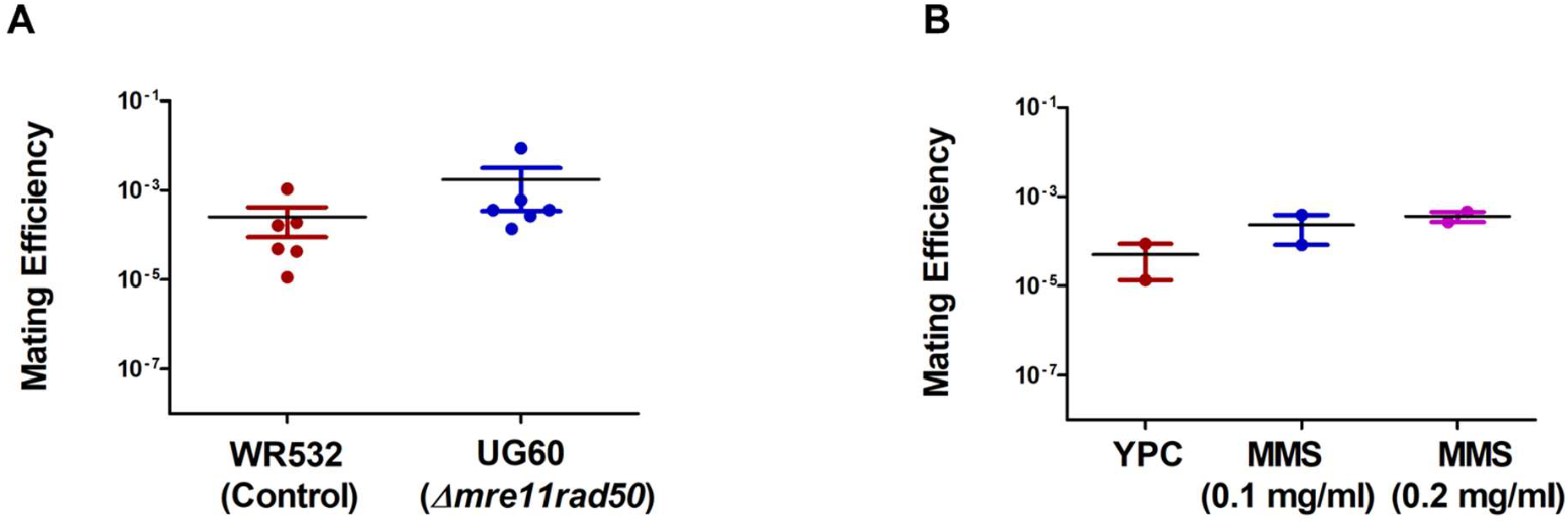
Disruption of *MRE11* and *RAD50* leads to increased mating efficiency in *H. volcanii*. **A**. Effect of *MRE11* and *RAD50 deletion* on archaeal mating frequency. The mating experiment was conducted between UG60 (*Δmre1, rad50, ΔpyrE)* and UG634 (*ΔhdrB, ΔtrpA*). Strain WR532 (*ΔpyrE*) was as control. The cells after mating were plated on Hv-Ca+ tryptophan. *MRE11* and *RAD50* deletion caused a significant increase in mating efficiency compared to the control. The Wilcoxon matched-pairs signed-rank test was used to compare control (WR532) and *Δmre1, rad50* (UG60) deletion mating, with a p-value of P < 0.0313 **B**. MMS treatment leads to increased mating efficiency in *H. volcanii*. The mating experiments were conducted between the UG532 (*ΔpyrE*) and UG634 (*ΔhdrB, ΔtrpA*) strains. YPC plates with and without MMS (0.1 mg/ml) and (0.2 mg/ml) were used. The cultures were incubated on YPC+MMS plates and YPC plates (used as a control) for 24 hours. The standard mating protocol was then followed. The results shown are an average of two independent experiments.

## Discussion

HGT is considered a crucial driver of microbial evolution. Previous studies have indicated that CRISPR-Cas systems limit HGT through mechanisms such as conjugation and plasmid transformation, due to their ability to defend against foreign DNA acquisition^20^. Previous work from our lab showed that CRISPR-Cas targeting can restrict gene exchange during inter-species mating between *H. volcanii* and *Haloferax mediterranei*^8^. However, the molecular mechanisms driving the interaction between CRISPR-Cas targeting and HGT within the same species have not been fully understood.

In this study, we demonstrate that CRISPR-Cas targeting significantly *increased* within-species mating efficiency in *H. volcanii*. This elevation of mating efficiency can be attributed to increased HR stimulated by CRISPR-Cas activity. Indeed, when we used a genetic background (mutated in *mre11* and *rad50*) known to have higher HR rates^9,19^. this resulted in higher mating efficiency. The direct association between higher HR and higher mating can also explain the seeming contradiction between the results presented here and our previous work, where CRISPR-Cas targeting *reduced* inter-species mating efficiency between genetically distant *H. volcanii* and *H. mediterranei*^8^. This discrepancy can be explained by the chances of successful HR being much higher when there is high sequence identity (within-species), as is long known for bacteria, eukaryotes and archaea^6,10^.

Mechanistically, the increase in mating and recombination frequency was probably stimulated by the breaks that Cas3 generates during CRISPR-Cas-mediated DNA cleavage. This increase was dependent on the presence of Cas3 and a targeting CRISPR spacer. In agreement with a CRISPR-Cas stimulated process, recombination favoured integration of genetic material from the genome that was cut into that of the mating partner doing the cutting. Mechanistically this is likely to be driven by a single strand of DNA from the lesion generated by Cas3 helicase-nuclease activity that invades the dsDNA of the CRISPR-positive mating partner.

The bias in recombination observed during targeting conditions is particularly intriguing. Our results show that CRISPR-Cas targeting leads to preferential gene flow from the targeted strain to the targeting strain. This genetic antagonism could provide a competitive advantage to strains possessing active CRISPR-Cas systems, as they can acquire beneficial genetic material from related strains while limiting the reciprocal exchange. This asymmetric gene flow could affect adaptation and evolution within archaeal populations.

The fact that DNA cleavage increased overall mating rates prompted us to investigate whether less-specific DNA damage will also increase mating frequency. Indeed, exposure of *H. volcanii* to the DNA alkylating agent MMS in cells with intact *mre11*-*rad50* also increased mating frequency, but only slightly. This is perhaps to be expected, since in *H. volcanii* break repair is done primarily by micro-homology mediated end-joining and not homologous recombination, in the presence of wild-type *mre11*-*rad50*^19,21^(Figure 4B). DNA repair by HR with an intact DNA template was also previously shown in the Sulfolobales, where UV exposure induced pili-mediated aggregation and subsequent genomic DNA exchange by the Ced system^22–24^, a process somewhat analogous to haloarchaeal mating, though different mechanistically.

In summary, the study presents strong evidence that CRISPR-Cas targeting boosts within-species mating efficiency in *H. volcanii* by enhancing HR rates. Since CRISPR-Cas targeting between different species was shown to have the opposite effect^8^, it therefore follows that CRISPR-Cas systems help maintain species barriers in halophilic archaea.

## Material and Methods

### Culture conditions

The wild-type *Haloferax* strains were routinely cultured at 45°C in either Hv-YPC or Hv-Ca medium. *H. volcanii* transformants were selected and grown in either Hv-Ca or Hv-Enhanced Ca (Hv-ECa) medium. Thymidine (40 μg/ml) and tryptophan (50 μg/ml) were added when needed. Bacterial strains were cultured at 37°C in LB medium, or in LB medium supplemented with ampicillin for strains carrying plasmids.

### Construction of *H. volcanii* strains with spacer+PAM sequence

The strain was constructed using the spacer+PAM sequence following the pop-in/pop-out protocol described in (Bitan-Banin, Ortenberg, and Mevarech 2003; Allers et al. 2004). In this method, the TrpA flanking regions (upstream with HindIII and ApaI restriction sites) and a 40 bp spacer+PAM sequence (ttcgcaggcatctcgaccggcgacctcccggaacactttg) along with the *TrpA* flanking downstream regions (upstream with ApaI and EcorI restriction sites) are amplified by PCR using specific primer sets. These amplified regions are then cloned into the non-replicating “suicide plasmid” pTA131, which harbors the *pyrE*2 selectable genetic marker.

The plasmids are transformed into *H. volcanii* mutant strains lacking cas genes, hdrB, and pyrE2. Transformants are selected on uracil-deficient media Hv-Ca (“pop-in”), where plasmids integrate into the chromosome. Subsequent counterselection on uracil and 5-FOA plates allows survival only of cells where integrated plasmids are excised through spontaneous intrachromosomal homologous recombination (“pop-out”). This process either restores the wild-type gene or achieves allele exchange. “Pop-out” strains are verified by PCR using primers flanking the deletion site and confirmed by Sanger sequencing. The parental strains and plasmids used in the constructions are detailed in Tables 1 and 2.

### Mating Protocol

The liquid cultures of both parental strains were grown separately until they reached stationary phase, as measured by optical density at 600 nm. The strains were then combined in a 1:1 ratio and filtered through 0.45 μm nitrocellulose filters using a Swinnex 25 mm filter holder. The filter containing the mating mixture was transferred to a rich medium plate (Hv-YPC supplemented with thymidine) and incubated for 48 hours at 45°C to allow for phenotypic expression. Following incubation, the cells were resuspended in Hv-Ca media and washed three times with the same media. Finally, the cell suspension was diluted and plated onto selective media chosen based on different mating markers.

### MMS treatment assay

The mating experiment was conducted between strains UG532 (*ΔpyrE*) and UG634 (*ΔhdrB, ΔtrpA*). The membrane-containing cultures were incubated on both YPC+MMS (0.1 and 0.2 mg/ml) and YPC-only plates for 24 hours at 45°C, after which the standard mating protocol was followed.

### Measuring Mating Efficiencies

Mating efficiency was calculated by dividing the average number of colony-forming units (CFU) on the selective mating plates by the average number of CFU of each parental strain grown on rich media plates.

## Supplementary Figures

**Supplementary Fig. 1.**
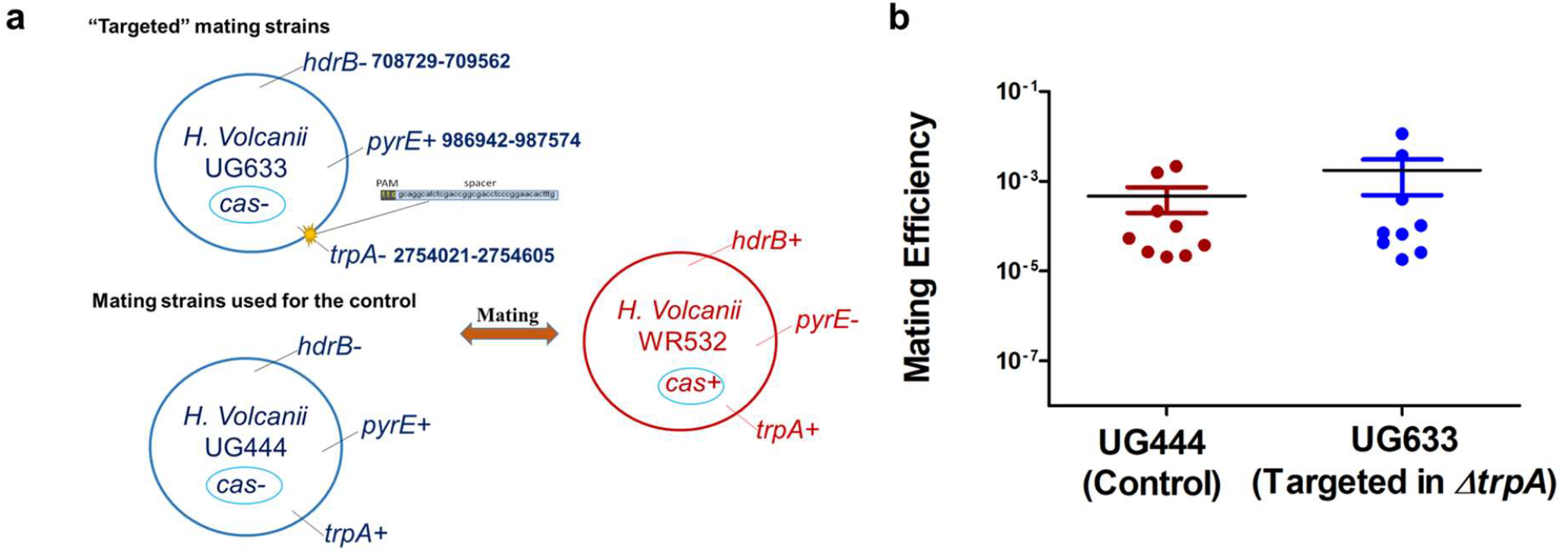
Increased mating efficiency was mostly due to CRISPR-Cas targeting, not the CRISPR-Cas machinery itself. **a)** Strains used for the mating experiments. Mating experiment was conducted using UG633 (*ΔhdrB, ΔtrpA, Δcas* genes, containing spacer inside *ΔtrpA*) and WR532 (*ΔpyrE2*), we used *H. volcanii* UG444 (*ΔhdrB, Δcas*) as a non-targeted control strain. **b)** The cells after mating were plated on Hv-Ca. The targeted strain exhibited increased mating efficiency compared to the non-targeted control. The Wilcoxon matched-pairs signed-rank test was performed to compare targeted and non-targeted mating with p-values of *P* < 0.0742.

**Supplementary Fig. 2.**
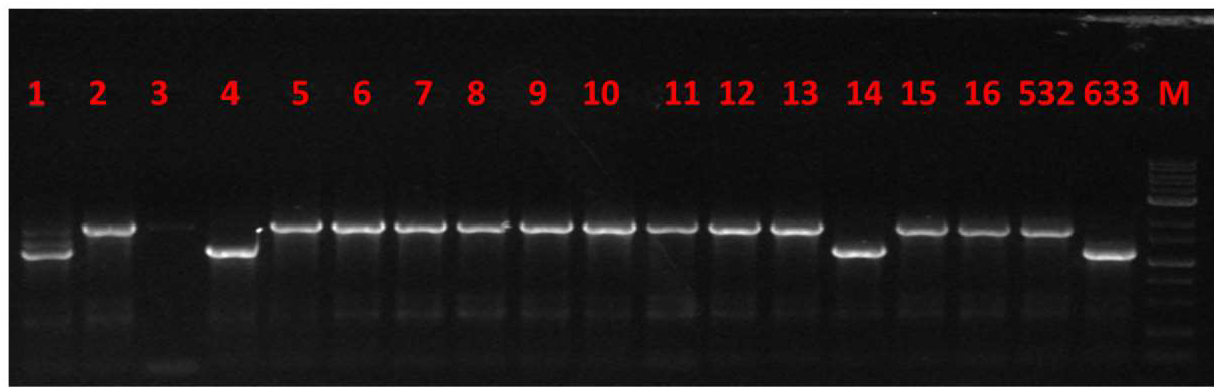
Representative PCR screening of the colonies to detect the trpA marker. The results of agarose gel electrophoresis, stained with ethidium bromide, of colony PCR products are shown. a) Colony PCR to check for the presence of the trpA marker after CRISPR-Cas targeting during mating. The gel displays colonies #1 to #16, from left to right, along with WR532 (ΔpyrE), UG633 (ΔhdrB, ΔtrpA, Δcas genes, contains spacer inside ΔtrpA), and a 1Kb DNA ladder (NEB). The expected smaller band size is ∼1200 bp, and the larger band size is ∼1692 bp.

**Supplementary table 1.**
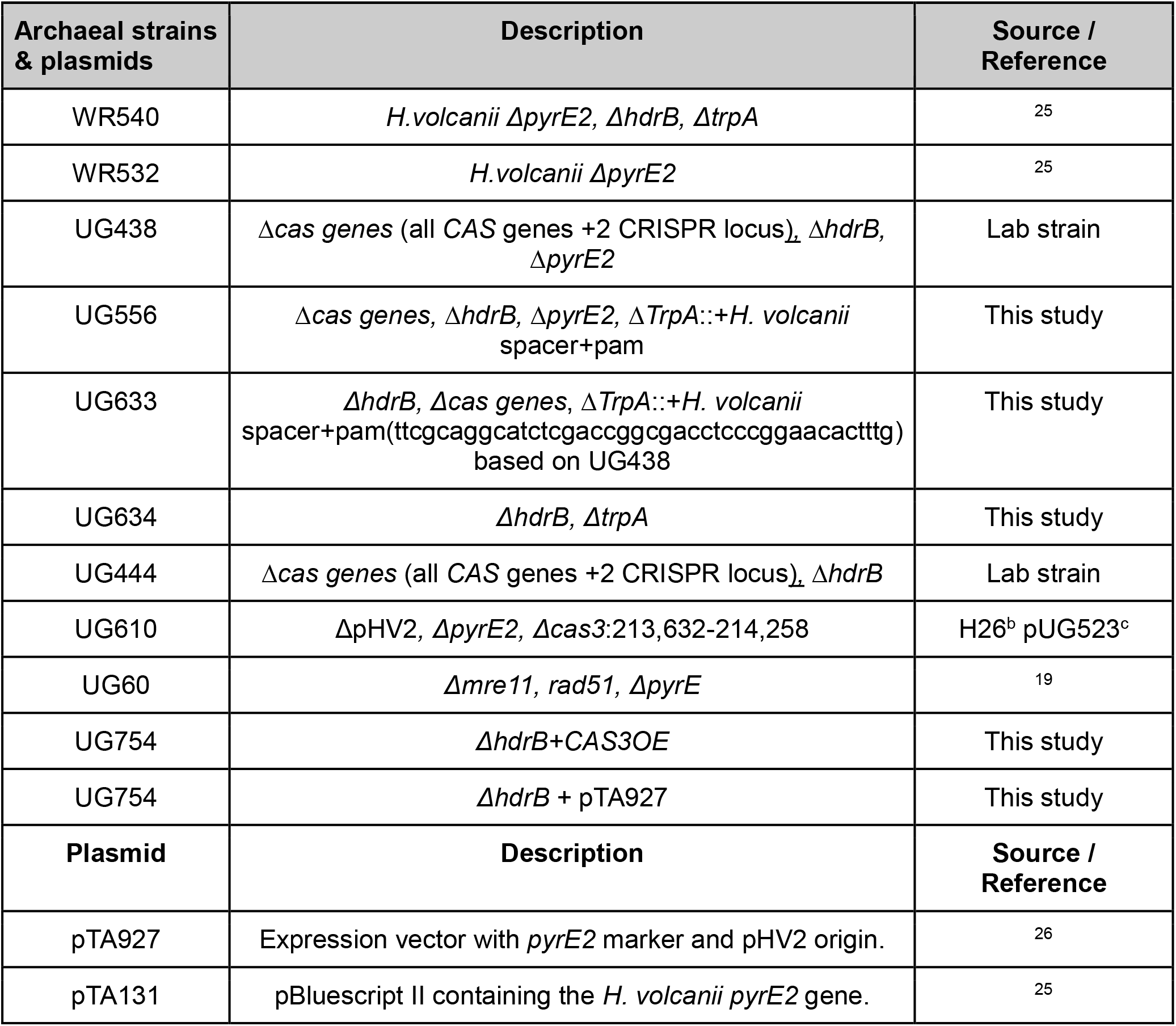
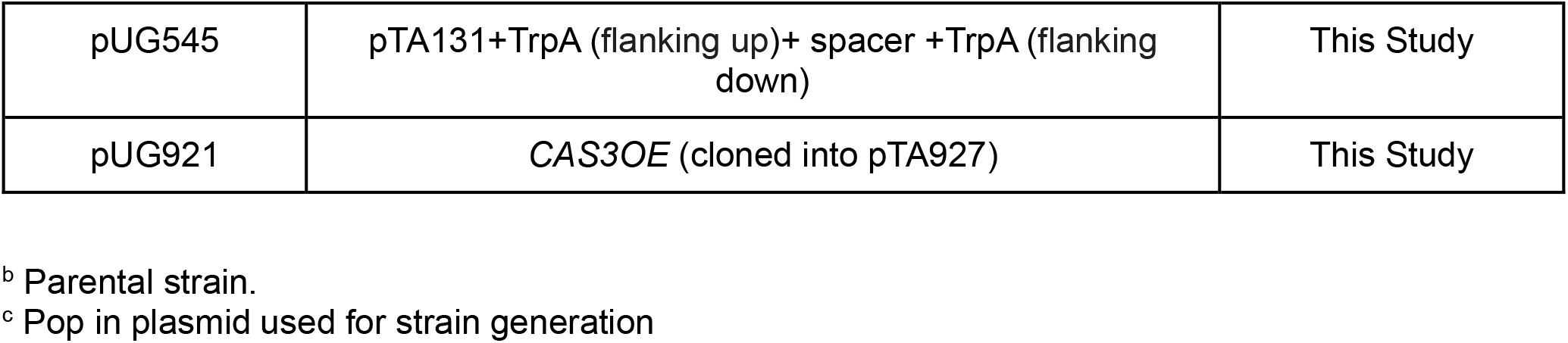
List of all the strains and plasmids used in this study.

**Supplementary table 2.**
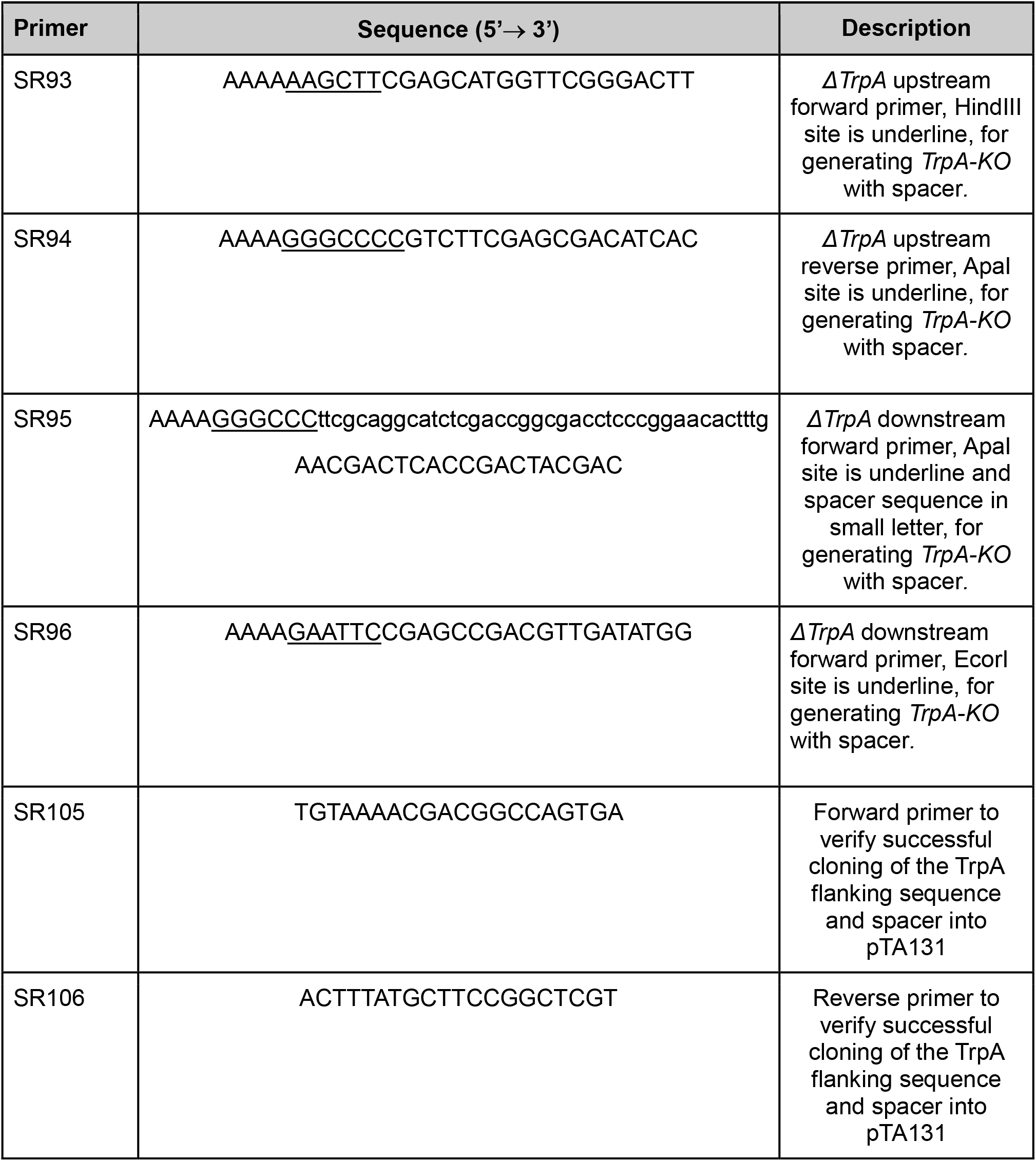
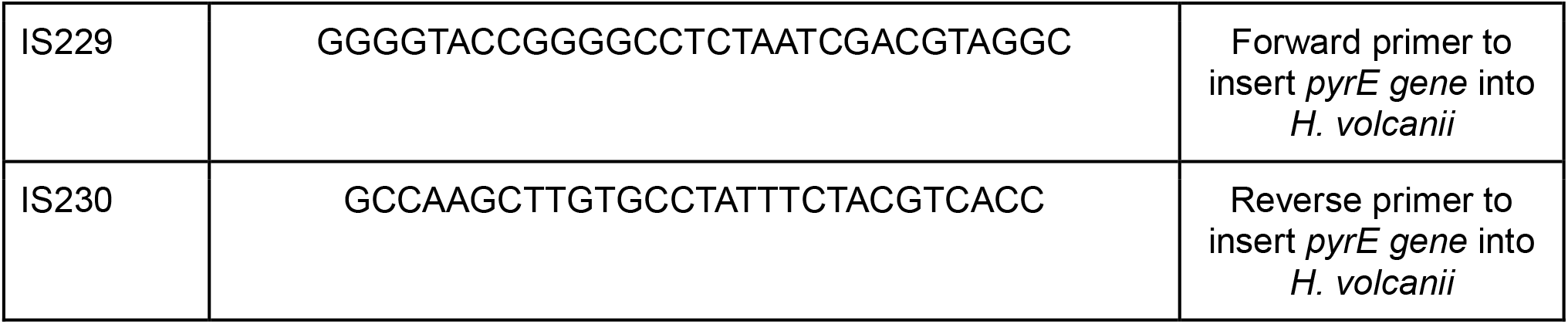
List of oligonucleotides used in this study.

## Author contributions

U.G., D.K.C. and I.T.G. conceived and designed the study. The CRISPR-Cas targeted strain was generated by S.B. and I.T.G. All the experiments were performed by D.K.C. D.K.C. prepared figures and analyzed the data with input from U.G. and I.T.G. The manuscript was written by D.K.C. and U.G., with input from I.T.G. All authors read and approved the draft.

## Acknowledgements

The authors thank Rimpa Mahanty for assistance with mating experiments. This work was funded by a European Research Council (grant ERC-AdG 787514) to UG and by the DFG priority programme “CRISPR-Cas functions beyond defence” SPP2141 grants to UG.

## References

1. Sorek, R., Kunin, V. & Hugenholtz, P. CRISPR — a widespread system that provides acquired resistance against phages in bacteria and archaea. Nat. Rev. Microbiol. 6, 181–186 (2008).

2. Makarova, K. S., Wolf, Y. I. & Koonin, E. V. Comparative genomics of defense systems in archaea and bacteria. Nucleic Acids Res. 41, 4360–4377 (2013).

3. McGinn, J. & Marraffini, L. A. Molecular mechanisms of CRISPR–Cas spacer acquisition. Nat. Rev. Microbiol. 17, 7–12 (2019).

4. Hille, F. & Charpentier, E. CRISPR-Cas: biology, mechanisms and relevance. Philos. Trans. R. Soc. B Biol. Sci. 371, 20150496 (2016).

5. Maier, L.-K. et al. The nuts and bolts of the Haloferax CRISPR-Cas system I-B. RNA Biol. 16, 469–480 (2018).

6. Naor, A., Lapierre, P., Mevarech, M., Papke, R. T. & Gophna, U. Low species barriers in halophilic archaea and the formation of recombinant hybrids. Curr. Biol. CB 22, 1444– 1448 (2012).

7. Naor, A. & Gophna, U. Cell fusion and hybrids in Archaea. Bioengineered 4, 126–129 (2013).

8. Turgeman-Grott, I. et al. Pervasive acquisition of CRISPR memory driven by inter-species mating of archaea can limit gene transfer and influence speciation. Nat. Microbiol. 4, 177–186 (2019).

9. Miezner, G. et al. An archaeal Cas3 protein facilitates rapid recovery from DNA damage. microLife 4, uqad007 (2023).

10. Cadillo-Quiroz, H. et al. Patterns of Gene Flow Define Species of Thermophilic Archaea. PLoS Biol. 10, e1001265 (2012).

11. Mevarech, M. & Werczberger, R. Genetic transfer in Halobacterium volcanii. J. Bacteriol. 162, 461–462 (1985).

12. Rosenshine, I. & Mevarech, M. The Kinetic of the Genetic Exchange Process in Halobacterium Volcanii Mating. in General and Applied Aspects of Halophilic Microorganisms (ed. Rodriguez-Valera, F.) 265–270 (Springer US, Boston, MA, 1991). doi:10.1007/978-1-4615-3730-4_32.

13. Fischer, S. et al. An archaeal immune system can detect multiple protospacer adjacent motifs (PAMs) to target invader DNA. J. Biol. Chem. 287, 33351–33363 (2012).

14. Wörtz, J. et al. Cas1 and Fen1 Display Equivalent Functions During Archaeal DNA Repair. Front. Microbiol. 13, (2022).

15. Sinkunas, T. et al. Cas3 is a single-stranded DNA nuclease and ATP-dependent helicase in the CRISPR/Cas immune system. EMBO J. 30, 1335–1342 (2011).

16. Gong, B. et al. Molecular insights into DNA interference by CRISPR-associated nuclease-helicase Cas3. Proc. Natl. Acad. Sci. U. S. A. 111, 16359–16364 (2014).

17. Brunner, E. et al. CRISPR-induced double-strand breaks trigger recombination between homologous chromosome arms. Life Sci. Alliance 2, e201800267 (2019).

18. Zhang, F., Cheng, D., Wang, S. & Zhu, J. Crispr/Cas9-mediated cleavages facilitate homologous recombination during genetic engineering of a large chromosomal region. Biotechnol. Bioeng. 117, 2816–2826 (2020).

19. Delmas, S., Shunburne, L., Ngo, H. P. & Allers, T. Mre11-Rad50 promotes rapid repair of DNA damage in the polyploid archaeon Haloferax volcanii by restraining homologous recombination. PLoS Genetics vol. 5 Preprint at 10.1371/journal.pgen.1000552 (2009).

20. Marraffini, L. A. & Sontheimer, E. J. CRISPR Interference Limits Horizontal Gene Transfer in Staphylococci by Targeting DNA. Science 322, 1843–1845 (2008).

21. Pérez-Arnaiz, P., Dattani, A., Smith, V. & Allers, T. Haloferax volcanii—a model archaeon for studying DNA replication and repair. Open Biol. 10, 200293 (2020).

22. Fröls, S. et al. UV-inducible cellular aggregation of the hyperthermophilic archaeon Sulfolobus solfataricus is mediated by pili formation. Mol. Microbiol. 70, 938–952 (2008).

23. Ajon, M. et al. UV-inducible DNA exchange in hyperthermophilic archaea mediated by type IV pili. Mol. Microbiol. 82, 807–817 (2011).

24. van Wolferen, M., Wagner, A., van der Does, C. & Albers, S.-V. The archaeal Ced system imports DNA. Proc. Natl. Acad. Sci. 113, 2496–2501 (2016).

25. Allers, T., Ngo, H. P., Mevarech, M. & Lloyd, R. G. Development of Additional Selectable Markers for the Halophilic Archaeon Haloferax volcanii Based on the leuB and trpA Genes. Applied and Environmental Microbiology vol. 70 943–953 Preprint at 10.1128/AEM.70.2.943-953.2004 (2004).

26. Allers, T. Overexpression and purification of halophilic proteins in Haloferax volcanii. Bioeng. Bugs 1, 288–290 (2010).

